# Highly Accurate and Precise Determination of Mouse Mass Using Computer Vision

**DOI:** 10.1101/2023.12.30.573718

**Authors:** Malachy Guzman, Brian Geuther, Gautam Sabnis, Vivek Kumar

**Affiliations:** The Jackson Laboratory, Bar Harbor, ME USA; Carleton College, Northfield, MN USA; School of Graduate Biomedical Sciences, Tufts University School of Medicine, Boston, MA USA; Graduate School of Biomedical Sciences and Engineering, University of Maine, Orono, ME USA

## Abstract

Changes in body mass are a key indicator of health and disease in humans and model organisms. Animal body mass is routinely monitored in husbandry and preclinical studies. In rodent studies, the current best method requires manually weighing the animal on a balance which has at least two consequences. First, direct handling of the animal induces stress and can have confounding effects on studies. Second, the acquired mass is static and not amenable to continuous assessment, and rapid mass changes can be missed. A noninvasive and continuous method of monitoring animal mass would have utility in multiple areas of biomedical research. Here, we test the feasibility of determining mouse body mass using video data. We combine computer vision methods with statistical modeling to demonstrate the feasibility of our approach. Our methods determine mouse mass with 4.8% error across highly genetically diverse mouse strains, with varied coat colors and mass. This error is low enough to replace manual weighing with image-based assessment in most mouse studies. We conclude that visual determination of rodent mass using video enables noninvasive and continuous monitoring and can improve animal welfare and preclinical studies.

## 2 Introduction

Body mass is a primary measure of health and disease in humans. For example, body mass index is a key measure of metabolic health, and is one of the oldest and most widely used metrics in modern medicine [1]. Changes in body mass indicate the function of many systems, including metabolic, cardiac, and the psychiatric, and are often the primary symptom of disease onset [2, 3]. Rodents, particularly mice, are commonly used to model human diseases, carry out preclinical studies, and investigate disease pathologies. Over 95% of disease research that involves animal models is conducted with mice [4]. As in humans, changes in body mass in mice are predictive of health, particularly when sudden changes occur [5]. The Institutional Animal Care and Use Committee (IACUC) includes significant body mass loss in its “humane intervention” guide, a set of standardized criteria that call for veterinary intervention or euthanasia of subjects. Loss of body mass greater than 20% relative to baseline or to matched controls is a common justification for such intervention [6].

Outside of ethical care, body mass is also an important feature collected during pre-clinical, metabolic, cardiovascular, and neurobiological studies [7]. However, handling mice to access mass induces physiological stress responses and alters immune responses [8]. Additionally, weight measurement is static; body mass is measured every few days during the animal’s rest phase depending on the protocol. Weight is often visually assessed during daily health checks by caretakers, and mass is measured only if the animal looks unhealthy or has lost weight. This can be subjective and can lead to variable treatment of the animal. These effects of human handling and static periodic measures of mass can weaken the validity and reproducibility of mouse experiments. Thus, there is a need for better methods to accurately and noninvasively measure animal mass over time in order to improve mouse models of disease and to improve animal welfare.

To address this problem, we explore a generalized solution to determine mass using computer vision. The application of computer vision to determine an animal’s body mass has been applied in industrial farming, including poultry farming, cattle, and fishing [9–11]. Visual mass determination in these animals is easier because these animals have relatively constant silhouettes because of their rigid bodies. Mice, on the other hand, are highly flexible, have much smaller but highly deformable bodies, and quickly change shape. This deformability is a direct result of animal behavior and is subject to individual differences in genetics. Thus, the deformability of rodents is a major challenge to visual weight assessment.

Other noninvasive methods to determine animal mass have been implemented using highly engineered cages with balances and compartments [5]. Although these methods alleviate animal handling issues and provide continuous mass measurement, the highly engineered cage designs require modification of housing conditions and are currently limited to singly housed animals. Thus, these methods can be challenging to scale and practically implement. A computer vision approach is suitable for multiple environments, both within and outside home cages, and can be scaled for continuous monitoring of mass in multiple mice. Therefore, this approach is potentially highly generalizable and scalable. However, whether this approach is accurate and precise enough to replace manual weight assessment, particularly in genetically diverse mouse strains, has yet to be determined.

Here, we test the use of single-camera video surveillance that functions autonomously and does not require the animal to take predetermined actions. We compare 6 statistical models with different use-cases on a dataset of 62 genetically diverse mouse strains. The best of these models predicts mass with a *R*^2^ of 0.92, mean absolute error (MAE) of 1.16 grams, and a mean absolute percentage error (MAPE) of 4.8%. This performance makes our approach applicable to preclinical studies, mouse husbandry, and other biological studies. It has the potential to expand weight assessment to novel environments and for continuous assessment. Additionally, this method can be easily incorporated into existing computer vision-based monitoring systems for routine use. We conclude that computer vision-based mass determination is highly accurate and precise and can be further developed for real-world applications.

## 3 Results

### 3.1 General Approach

Our approach to visual mass determination is outlined in Figure 1A and detailed below. Briefly, we collect video data from the top-down perspective in an open field [12]. Each frame is segmented for the mouse, which describes the animal’s size. The segmented image is used to fit an ellipse that describes the approximate posture of the animal and is then adjusted with covariates to improve our modeling and mass prediction.

**Figure 1:**
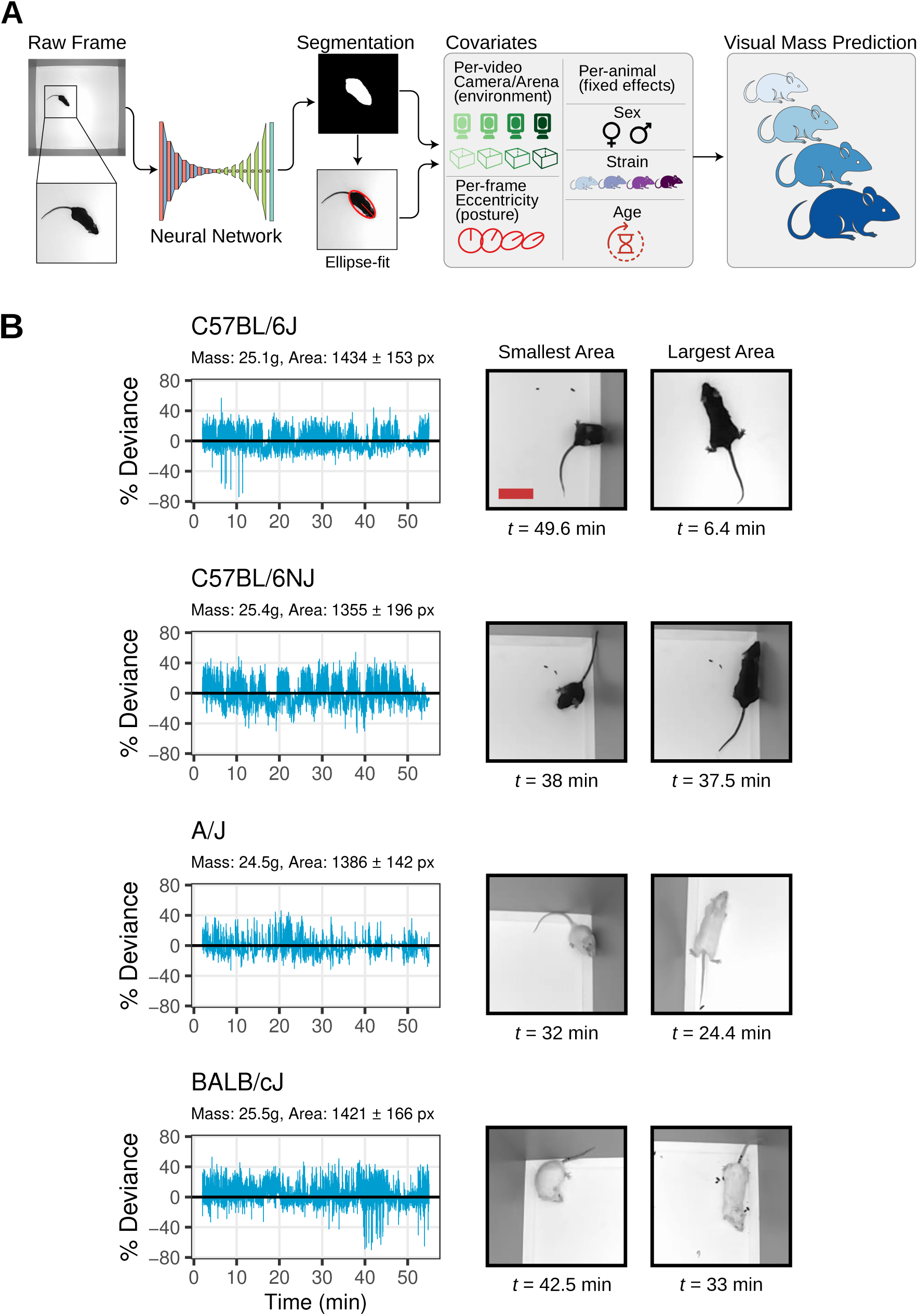
Visual mass determination approach taken to address highly variable segmentation areas observed in our data. (A) Flow chart describing the full computational process from open field video to body mass prediction. (B) Time series of percent deviations from the mean segmentation area over 55 minutes for four individual C57BL/6J, C57BL/6NJ, A/J, and BALB/cJ mice, with (mean ± SD) of pixel area reported. Raw frames of the approximate least and greatest segmentation areas are shown to the right. Red bar on the C57BL/6J smallest frame indicates 5 cm.

Mice are highly deformable, depending on their behavior. To determine the effect of deformability on the segmentation area, we inspected the variation of segmentation area in 55-minute videos of several mice that are approximately the same mass (25.1g) from different strains Figure 1B. We found that the area of the mouse commonly deviated by ±40% (relative to mean area) over the course of the video (Figure 1B, Left). Upon examining the video frames, we saw that the high variation in area was due to deformability of the animal due to altered behavior (Figure 1B, Right). Over a short time frame, the mouse scrunches, stretches, bends, and rears such that its segmentation area quickly changes.

Since we have multiple images for each animal we focus on the mean or median segmentation area. However, variance in this measure can impact the precision of a prediction. We investigated this by exploring changes in variation of segmentation area between these mice. We found that this variation is strain-specific, i.e. dependent on genetics. For instance, a 25.4g C57BL/6NJ mouse has an average area of 1355 ± 196 px, whereas a smaller 24.5g, A/J mouse has a larger area of 1386 ± 142 px. However, the variation in area is larger in C57BL/6NJ than in A/J. A/J are a high anxiety strain with low ambulation, while C57BL6/NJ have more bouts of activity [13, 14]. The larger segmentation area and lower variation in A/J is because it spends more time in the corner in a constant posture. We reasoned that simple segmentation is inadequate to handle genetic diversity seen in laboratory mice, and a more sophisticated approach is needed.

### 3.2 Data and Data Acquisition

Our dataset consists of 2,028 videos obtained from a previously conducted strain survey experiment [12]. For an example of our video data, see Supplementary Video 1, where we provide ten-second clips from the videos corresponding to the frames in Figure 1B. Sampled mice come from 62 different strains, with most strains represented by at least 4 male and 4 female mice (Table S1). Most are 8-12 weeks old, but vary from 6-26 weeks old, with a mean of 11 weeks old. Body mass was measured immediately before open field recording with a precision laboratory scale, and ranges from 9.1 g - 54 g, with a mean of 24.7g. Each video has one mouse in an open field enclosure which measures 52 *×* 52 cm. Videos are 480 *×* 480 px, were recorded at 30 FPS, last 55 minutes, and have 8-bit monochrome depth. The camera is positioned approximately 36 inches (91.44 cm) above the floor. A 25.1g C57BL/6J mouse occupies 1434 pixels on average, or 0.6% of the total image pixels (Figure 1A,B) [15]. This is similar to the distance and mouse pixel area if a camera were placed on top of a mouse cage changing table used in most barrier mouse facilities.

### 3.3 Error Metrics

To evaluate fit accuracy, we used the coefficient of determination *R*^2^. To evaluate model error, we used Mean Absolute Error (MAE), Root Mean Square Error (RMSE), and Mean Absolute Percentage Error (MAPE) (see Methods for detail) [16–18]. We used MAE to get an intuitive sense of average error, RMSE to penalize larger errors more strictly, and MAPE to get error relative to mass. Each of these metrics are commonly used to evaluate performance of similar computer vision solutions.

### 3.4 Per frame segmentation

First, we process a raw video from the dataset through one deep neural network to predict a segmentation mask for the mouse for every frame of the video (Figure 1A). The segmentation network has been trained on a diversity of mouse images and achieves high accuracy [12]. We fit an ellipse to the segmented blob as an approximation of mouse posture.

In every video, we compute the pixel area of the segmented mouse image in each frame by summing the number of “mouse” pixels, which are identified by the segmentation mask. The white pixels in the “Segmentation” frame of Figure 1A are an example of mouse pixels. We summarize each video by the median of the pixel areas across all frames in the video, denoted *A_px_*. This provides a single area measurement per-mouse per-video that we can use for modeling.

### 3.5 Per Camera and Arena Correction

We collect data in 24 open field arenas. Although we have standardized video data collection to a tight tolerance, there is still slight variation in image acquisition between arenas. To account for differences in camera lens zoom, we converted the measured segmentation (square-pixel units) of the videos to real-world metric units (square cm units). Since all our arenas are manufactured to measure 52cm *×* 52cm, we located each arena’s corners in each video by applying a corner detection deep neural network, thereby deriving a scaling factor from pixels to centimeters [19]. To account for arena variance, we adjust the segmentation measurement *A_px_* by the following equation: *A_cm_* = *A_px_*/(L*_px_*/52cm)^2^, where *A_cm_* is area in square cm, *A_px_* is area in square pixels, and L*_px_* is corner edge length in pixels. This adjustment does not necessarily require knowledge of which arena the mouse was measured, one only needs to know the real world distance between corners, which should be a constant between all arenas in an experiment. If an experiment uses arenas of intentionally varying size, then *A_cm_*’s equation would have an arena-based variable instead of 52cm.

### 3.6 Posture Correction

So far, we have identified a metric, *A_cm_*, which captures the segmentation area of a single mouse, normalized for differences in camera zoom. However, this metric does not account for the wide variation in area for individual mice over short time periods observed in Figure 1B, which we determined was a direct result of changing behavior and posture in the mice. We reasoned that accounting for varying posture is necessary to increase performance of our visual method.

To correct for posture, we examined the effect of normalizing the area based on geometric shape descriptors of the segmentation mask. We tested several geometric shape descriptors, including eccentricity, aspect ratio, and elongation, as well as the speed of the mouse, as defined and evaluated in Table 1. We tested these shape descriptors under the assumption that changes in area in a short time frame are not related to changes in mass, and so must be related to changes in pose. To determine which shape descriptor works best, we compared RMSE, MAE, MAPE, and *R*^2^ values for each of four models (T1-T4, Table 1). Each model is a single variable linear regression, taking the product of *A_cm_* and the median of one posture metric (eccentricity, aspect ratio, elongation, or speed) as its input variable and predicting body mass. These models are evaluated under 50-fold cross-validation with a 70-30 training-testing split, explained in full detail in Section 5.3, Statistical Analysis.

**Table 1:** Posture Adjustment Metrics. Models T1-T4 have one input variable each, the product of *A_cm_* and the median of the given metric. T1’s input is equivalent to *A_e_* in the previous section, and the inputs of T2-T4 are analogous to *A_e_* but for aspect ratio, elongation, and speed. In the Metric Formulas, w and l are the width and length of the fitted ellipse, v*_x_* and v*_y_* are the x and y components of velocity (also representable as 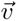), and each µ*_ij_* is a central image moment, calculated with the python package `OpenCV`.

| Model | Input Variable | Metric Formula | RMSE (g) | MAE (g) | MAPE (%) | R<sup>2</sup> |
| --- | --- | --- | --- | --- | --- | --- |
| T1 | $A_{cm} \times Med(eccentricity)$ | $(w^2 + l^2)^{1/2}/l$ | $2.353 \pm 0.054$ | $1.759 \pm 0.038$ | $7.328 \pm 0.173$ | $0.827 \pm 0.01$ |
| T2 | $A_{cm} \times Med(aspect\ ratio)$ | $w/l$ | $3.569 \pm 0.081$ | $2.830 \pm 0.06$ | $12.298 \pm 0.275$ | $0.603 \pm 0.022$ |
| T3 | $A_{cm} \times Med(elongation)$ | $\frac{\mu_{20} + \mu_{02} + \sqrt{4(\mu_{11})^2 + (\mu_{20} - \mu_{02})^2}}{\mu_{20} + \mu_{02} - \sqrt{4(\mu_{11})^2 + (\mu_{20} - \mu_{02})^2}}$ | $16.83 \pm 15.77$ | $4.714 \pm 0.414$ | $21.059 \pm 2.196$ | $0.170 \pm 0.125$ |
| T4 | $A_{cm} \times Med(speed)$ | $\sqrt{(v_x)^2 + (v_y)^2}$ or $\vert \vec{v}\vert $ | $5.644 \pm 0.127$ | $4.422 \pm 0.095$ | $19.487 \pm 0.460$ | $0.002 \pm 0.003$ |

Model T1 uses hyperbolic eccentricity—defined as e = (w^2^ + l^2^)^1^*^/^*^2^/l, where w, l are width and length—as its shape descriptor, T2 uses aspect ratio, T3 uses elongation, and T4 uses speed, all defined in Table 1. With hyperbolic eccentricity, T1 achieves an *R*^2^ of 0.827 and MAPE of 7.328%, significantly better performance than models T2-T4. This led us to conclude that eccentricity is the most useful indicator of shape of the metrics we tested. We initially hypothesized that speed would be a good descriptor to correct posture by, since mice tend to take on a constant elliptic shape when they’re moving. However, speed performs poorly, resulting in a MAPE of 19.4% and *R*^2^ value of only 0.002. We hypothesize that this lack of quality is due to the proportion of time mice actually move quickly. Although mice do take on relatively constant postures while walking forward, this kind of motion takes up a relatively tiny fraction of total frames, and is likely strain specific. Thus, speed may work for certain strains, but does not generalize well in genetically diverse populations.

From this result, we formalize the “eccentric area” metric as *A_e_* = *A_cm_ ∗* e, where *A_cm_* is the median of unit-converted area and e is the median eccentricity, both over all frames of the given video. In full, *A_e_* = (*A_px_*/(L*_px_*/52cm)^2^) *∗* (w^2^ + l^2^)^1^*^/^*^2^/l. *A_e_* functions as our video-level summary metric. To quantify the effect of correcting posture with eccentricity, we compared the variation in *A_cm_* (anecdotally shown to be high in Figure 1B) and *A_e_* by computing their respective Relative Standard Deviations (RSD) across all strains (Figure 2, *A_cm_* in blue vs. *A_e_* in red). Note that these RSDs are of the observed area values, and are not necessarily comparable to RSDs of model prediction (see Supplementary Material for more detail).

**Figure 2:**
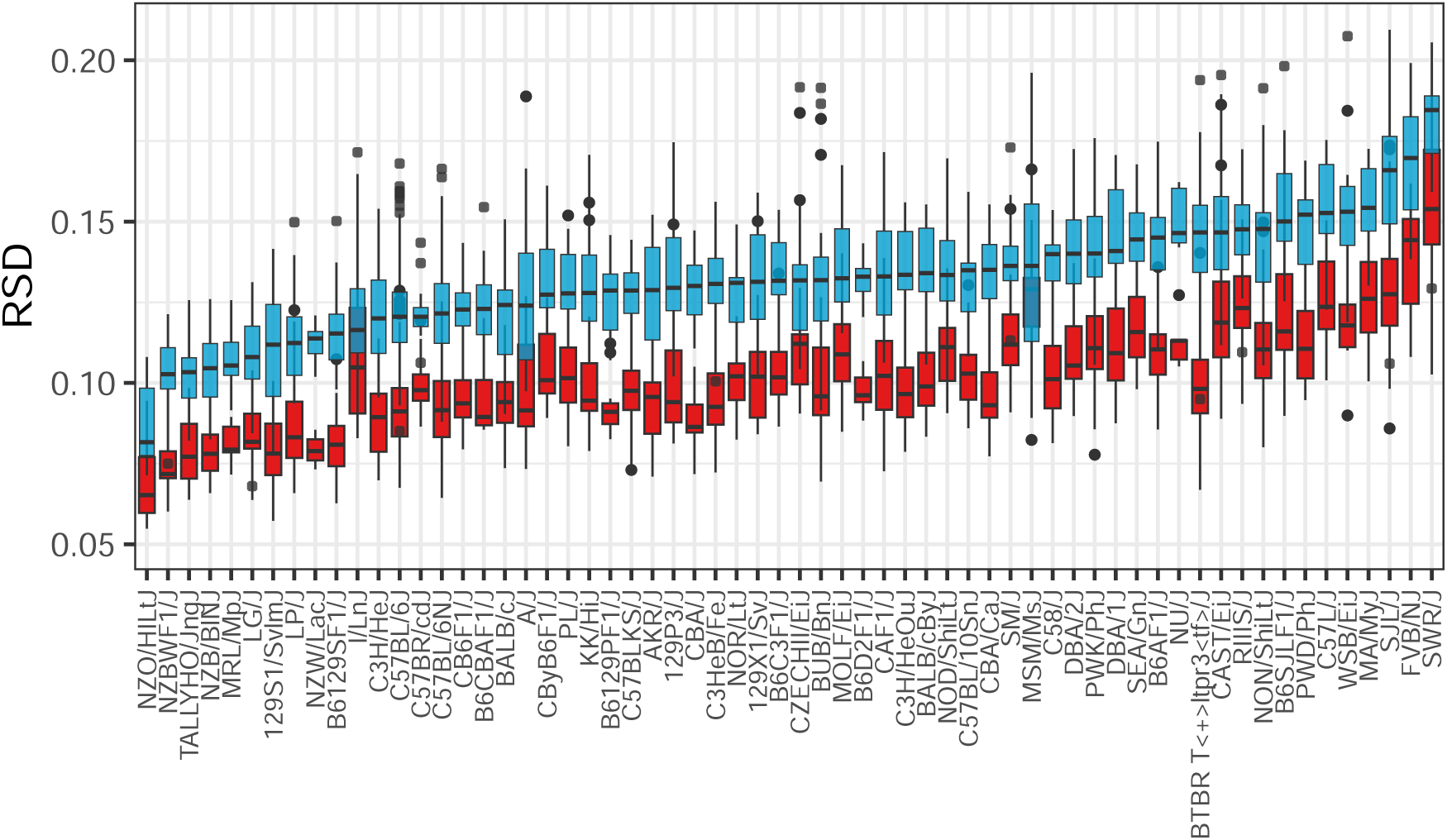
Adjustment of Segmentation Area by Eccentricity Reduces Relative Standard Deviation Across Strains. Relative standard deviations (RSD) of segmentation area *A_cm_* (in blue) and the adjusted size *A_e_* (in red) for each of the 62 mouse strains considered in our dataset.

Strain is another important factor in variability, as some strains like the obesity model NZO/HILtJ have a low RSD of around 6%, whereas a wild-derived strain like WSB/EiJ has a RSD around 15%. The largest area RSD we observed was about 18%, for the SWR/J strain. We hypothesize that these strain-level differences in RSD are largely due to differing activity levels and behavioral patterns between strains. After posture correction relative standard deviation across all strains is significantly reduced (Figure 2, in red). The RSD, however, is still strain dependent, indicating there here are still strain level effects on the precision of our prediction even after posture correction.

### 3.7 Other Covariate Correction

Equipped with several ways of describing size (*A_px_*, *A_cm_*, and *A_e_*), we built six multiple linear regression models (M1-6) to predict body mass. Each model uses one of raw area (*A_px_*), unit-converted area (*A_cm_*), or eccentric area (*A_e_*) as its visual variable, and different subsets of sex, strain, age, and arena (the specific open field the mouse was measured in) as covariates (Figure 3A-D). After building each model, we performed a 50-fold cross validation on a 70/30 training/testing split. Averaging our reported accuracy and error values over these 50 iterations ensures our analysis is not biased by lucky or unlucky sampling.

**Figure 3:**
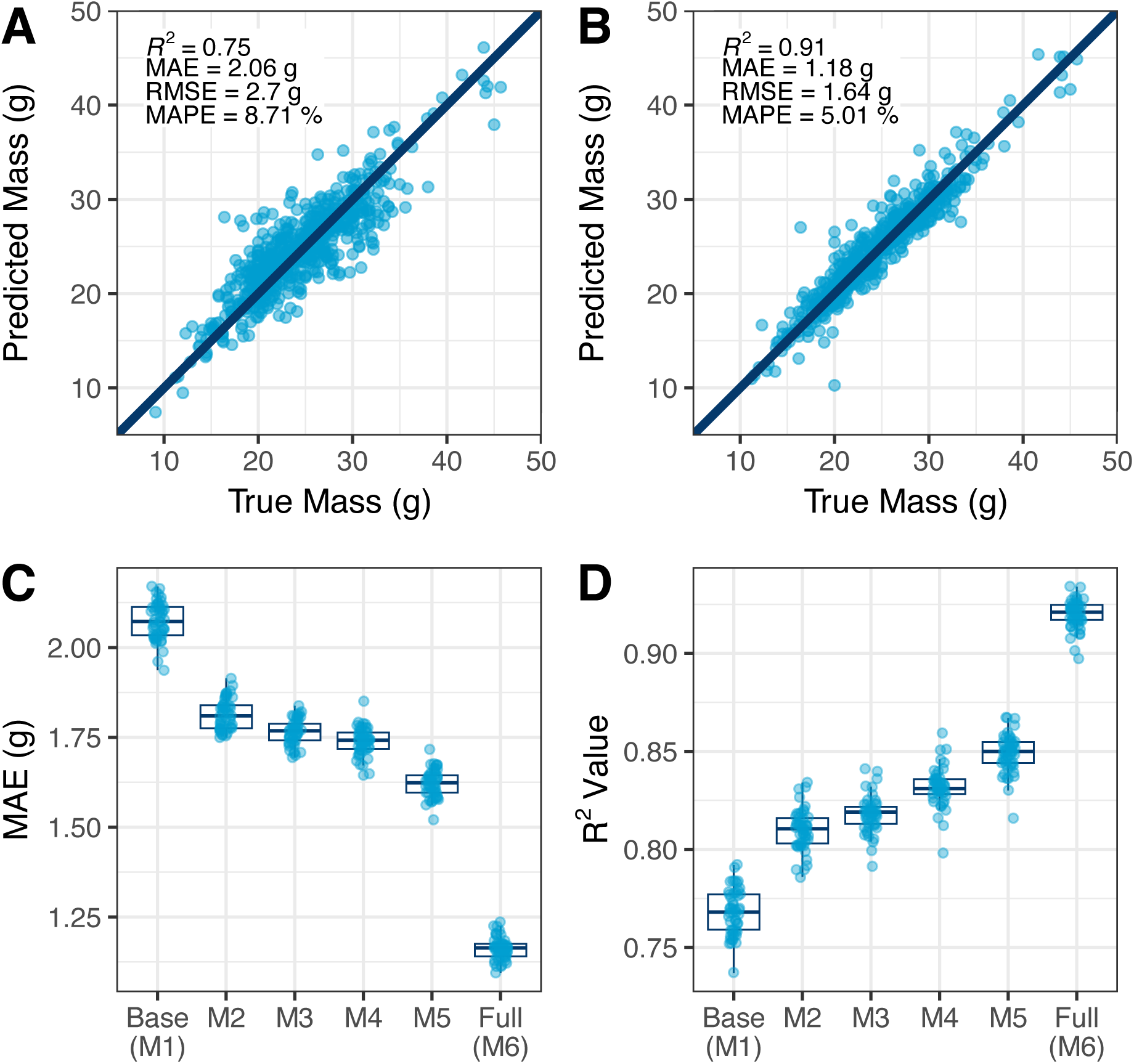
Model Performance. (A and B) Comparison of True Mass, as measured with a regular scale, and Predicted Mass, as determined by our process. Data shown is a one test sample. *R*^2^, Mean Absolute Error (MAE), Root Mean Squared Error (RMSE), and Mean Absolute Percentage Error (MAPE) presented. The 45*^◦^* line indicates perfect prediction. (A) Prediction quality for the Base Model (M1) accounting for only segmentation area. (B) Prediction quality for the Full Model (M6), accounting for true size *A_e_*, arena, sex, age, and strain. (C and D) Error and accuracy performance for each model under 50-fold cross-validation. (C) MAE across models. (D) *R*^2^ value across models.

Our first model, Base (M1), is a single variable linear regression that uses raw segmentation area *A_px_* to predict body mass, which we used as a comparative baseline for the other models (Table 2). This model has a *R*^2^ value of 0.767 and MAPE of 8.584% (Figure 3A). The second model, M2, is also a single variable regression, but between unit converted area *A_cm_* and body mass. M2 performs better than the Base model, increasing accuracy and decreasing error. In M3 we added a second variable, arena, the particular arena a mouse was tested in. M3 performs slightly better than M2 by all measures. M4 introduces eccentricity, which is combined with unit converted area to be one variable *A_e_*, as previously described; M4 has two variables: *A_e_* and arena. M4 performs substantially better, with *R*^2^ of 0.822 and MAPE of 7.566%.

**Table 2:** Model Definitions and Performance.

| Model | Model Input Variables | RMSE (g) | MAE (g) | MAPE (%) | $R^2$ |
| --- | --- | --- | --- | --- | --- |
| Base (M1) | $A_{px} (px^2)$ | $2.729 \pm 0.070$ | $2.072 \pm 0.052$ | $8.584 \pm 0.210$ | $0.767 \pm 0.012$ |
| M2 | $A_{cm} (cm^2)$ | $2.469 \pm 0.058$ | $1.814 \pm 0.042$ | $7.528 \pm 0.179$ | $0.810 \pm 0.010$ |
| M3 | $A_{cm} (cm^2)$, arena | $2.416 \pm 0.052$ | $1.765 \pm 0.035$ | $7.314 \pm 0.156$ | $0.818 \pm 0.009$ |
| Geometric (M4) | $A_e (cm^2)$, arena | $2.322 \pm 0.052$ | $1.739 \pm 0.039$ | $7.223 \pm 0.174$ | $0.832 \pm 0.010$ |
| Sex and Non-Genetic (M5) | $A_e (cm^2)$, arena, sex, age | $2.201 \pm 0.065$ | $1.622 \pm 0.037$ | $6.774 \pm 0.167$ | $0.849 \pm 0.010$ |
| Full (M6) | $A_e (cm^2)$, arena, sex, age, strain | $1.606 \pm 0.069$ | $1.162 \pm 0.031$ | $4.836 \pm 0.137$ | $0.920 \pm 0.007$ |

In a practical case of assuming minimal information, one would probably use M4 instead of M1, M2, or M3, because *A_px_*, *A_cm_*, and *A_e_* are visual metrics, and the particular arena is inherent to the experiment. We could always measure these variables and would have no reason not to, thus not using M4 in this case would decrease prediction performance for no experimental benefit. However, we present each model here because selectively adding and/or modifying variables demonstrates the performance benefit of each of unit conversion, arena identity, and eccentricity without confounding the variables. Additionally, likelihood-ratio tests confirm each model performs significantly better than the last, so one could use any model with confidence.

In many experiments the sex and age of the animal is known. Therefore, our fifth model, Sex and Non-Genetic (M5), adds sex and age as separate variables, reflecting the differences in body composition between sexes and as a mouse ages. This again provides a boost in accuracy and reduction in error values. In certain conditions, the strain of the mouse is also known. Therefore, the sixth model, Full (M6), introduces the strain of the mouse as a variable. Here, we include an interaction term between strain and sex, because we observed that while males are generally heavier than females, there are some strains for which the opposite is true. The Full model performs much better than the previous models M1-M5, achieving a mean *R*^2^ of 0.92 and MAPE of 4.84% (Figure 3B). Visualizing MAE and *R*^2^ values of each model illustrates their relative improvements under cross-validation (Figure 3C-D). The performance values for each model are compared in Table 2.

### 3.8 Models and Performance

### 3.9 Effect of Sex and Genetic Diversity

We observe large differences in mean body mass between sexes due to underlying changes in body composition. For most but not all mouse strains, males have higher percent fat than females [20]. We tested the performance of the full model (M6) on males and females. While females weigh less than males on average, our model performs uniformly well across females and males (Figure 4A). Differences in performance are slight: females and males have MAPE of 4.82 and 5.17%, respectively.

**Figure 4:**
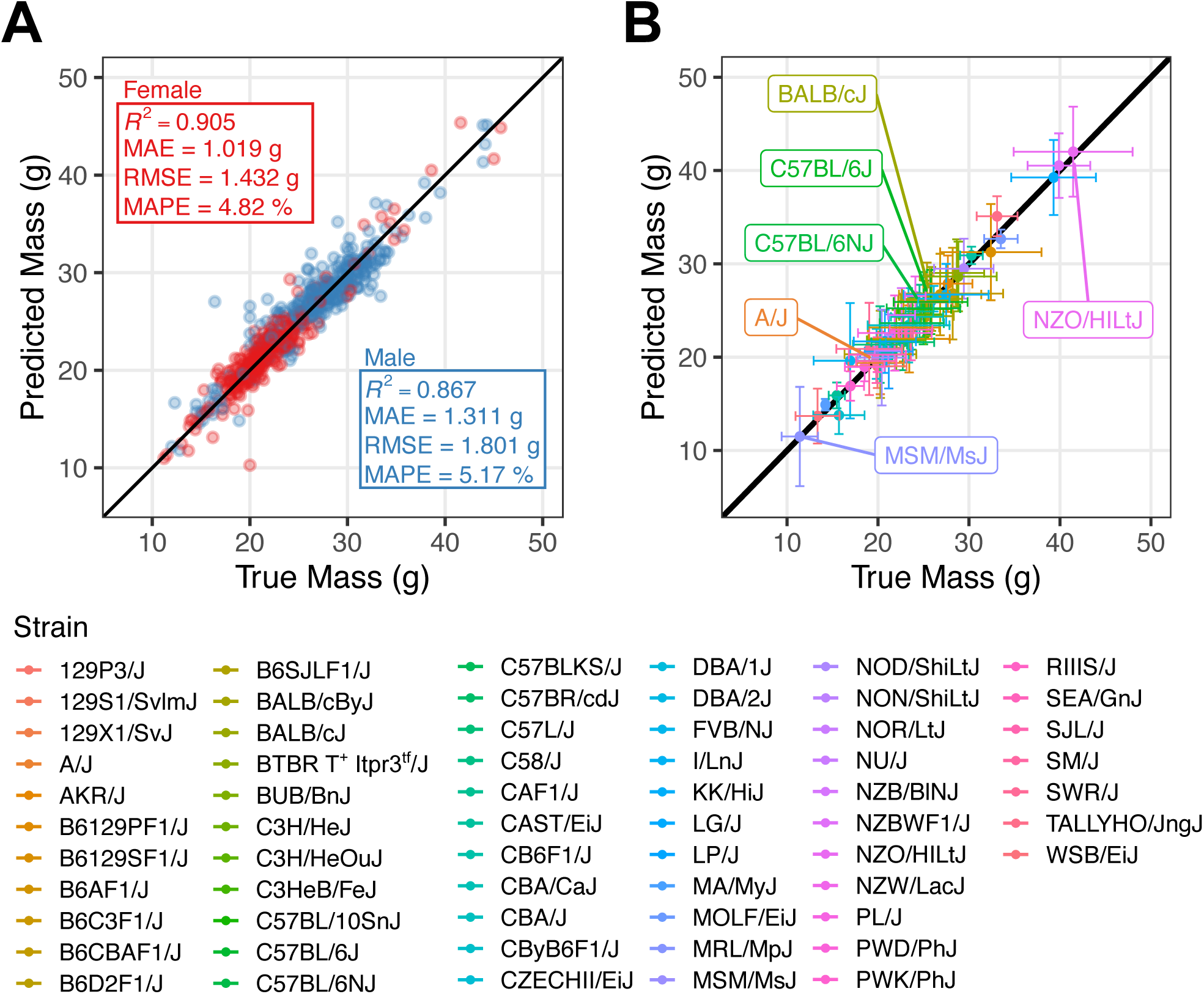
Full Model (M6) Performance by Sex and Strain. (A) Full model performance faceted by sex, with *R*^2^, MAE, RMSE, and MAPE values presented for males (in blue) and females (in red). (B) Full model performance faceted by strain (mean ± standard deviation). The four strains from Figure 1B are labeled here, along with the strains with least and greatest mean area. Test and train data is the same as in Figure 3B, for ease of comparison. Full results are presented in Table S1

Next, we tested performance of the full model (M6) across genetically diverse mice and measured accuracy and precision of the model across 44 classical inbred, 7 wild-derived inbred, and 11 F1 hybrid strains (Figure 4B). Most strains are between 20-30 g with wild-derived strains like MSM/MsJ at the low end and obesity models such as NZO/HILtJ on the high end of the distribution (labeled in Figure 4B). The model accurately predicted mean and variance across highly genetically diverse mice (Figure 4B, Figure S1, Figure S2). The variance of predicted mass reflects the variance of true mass, and suggests that the model captures much but not all variation within strains (Figure S1, Figure S2). We quantify the mean observed and predicted mass and standard deviation for each strain in Table S1. Additionally, we retrain model M6 on our entire dataset, and present the observed and predicted relative standard deviation in mass for each strain, demonstrating that real strain-level variation is captured in our model (Figure S1 and Figure S2). This final model can be used for inferring on any new data. We conclude that our model performs well across sex and diverse genetics.

## 4 Discussion

Body mass is an important measure of health in mice, and is widely used as both a feature of veterinary intervention and of behavioral and drug trial experiments. However, frequently monitoring body mass by hand is impractical over a long time period, and more importantly can negatively affect the actual experiment being performed. We sought to test whether visual methods provide adequate accuracy and precision to be a viable alternative to manual weighing of animals. Mice are highly deformable and change their posture in very short timescales. In addition, our camera is far from the mouse and the animal only 0.6% of the pixels in the video frame belong to the mouse. Thus, we do not have a very high resolution view of a highly deformable object. To solve this problem, we applied computer vision methods to develop models that predict the body mass of an individual mouse in an open field arena. We developed six models for this, each using a selection of visual information extracted from the segmentation and ellipse-fit masks, the particular arena a mouse was recorded in, and biological attributes of the mouse such as age, sex, and strain.

There are several advantages to our approach. First, our approach uses an explainable machine learning approach. The first step of our approach is to generate segmentation masks for the mouse. Segmentation is highly flexible and is one of the oldest and most established computer vision tasks particularly for biomedical images [21, 22]. Modern methods of segmentation improve upon traditional methods by using neural networks [23–27]. Segmentation neural networks can be trained with using much lower training data, and there are already pretrained models that for general purpose segmentation. For instance, we trained a segmentation network for sleep state prediction using 313 frames [28]. We selected a segmentation step to enable observable detection for accuracy shifts. For example, if a mouse escapes the arena, we can detect that no segmentation is predicted and make no body mass prediction. After segmentation, we apply well understood and explainable techniques including unit transformations, normalization, and linear regression modeling. Thus, our method is flexible for adoption to new environments and organisms without the burden of creating large training data while producing explainable results.

A second advantage of our approach is the use of highly diverse mouse strains, with varied coat colors, sizes, and behaviors, for training our models. This ensures that the model can handle diverse visual and size distributions that may be seen in real world settings. Each of our six models have different experimental applications. Though it is clear the Full model has the highest accuracy and lowest error, it is not necessarily suitable for all experiments. Practically, one would apply the Geometric, Non-Genetic, or Full model to an experiment, not M1, M2, or M3. While in some sense M1 is the most general of all the models, there’s no information in the Geometric model (M4) that is not easily attainable. Its two variables, *A_e_* and arena, are derived from the same base information in the video, not from any further information about the mouse itself. For the same reasons, it would equally make sense to use M4 instead of M2 or M3. As mentioned in the previous section, it is still useful to present M1-M3, because they show the benefit of additions and modifications to the models.

With fine-tuning to the particular environment, the Geometric model (M4) can be applied to all open field videos of similar size and resolution with an individual mouse. However, it would be a rare mouse model experiment that intentionally did not take into account the sex and age of the mice, as these are often important variables. The Sex and Non-Genetic model (M5), as its name suggests, would be very useful for experiments where collecting strain information is unfeasible or undesirable. A key example of this are experiments which use genetically diverse populations of mice, such as Diversity Outbred (DO) or Collaborative Cross (CC) populations. These types of populations are widely used to model genetic diversity, opposed to the standard inbred mouse strains, for example with application to high-resolution quantitative trait locus mapping (QTL), drug trials, drug exposure experiments, and disease experiments. In these cases, it is not necessarily easy to determine the genetics of a given mouse, and genetic information may give little insight in the context of body mass prediction anyway, since the mouse may be unique or one of only a handful in the particular screen. This means our strain-free model (M5) is highly applicable to diverse mouse model experiments. The Full model, which includes genetic information, is readily applicable to any single-mouse open field experiment where strain is relevant and known. Examples include mouse colony management, where strain is known and highly relevant to body composition. Our Full model would provide the highest accuracy monitoring of body mass in these areas. Furthermore, any classical mouse experiment with inbred strains that lends itself to having one mouse in an open-field environment would benefit from this model, especially behavioral experiments where inducing anxiety potentially introduces bias.

For health monitoring, this style of visual prediction is only useful if its error falls well within the standards of humane intervention points, as to avoid false negative or false positive determinations of medically concerning body mass fluctuation. Following Institutional Animal Care and Use Committee (IACUC) Humane Intervention guidelines, it is necessary to intervene, typically with euthanasia, if a mouse loses more than 20% of its body mass compared to similar control animals. As an example, if an inbred mouse lost 20% of its body mass during an experiment, our results indicate we could use the Full model to predict that it had lost approximately *∼*15.2-24.8% of its body mass, which is a small enough range that we know something is likely wrong with the mouse, and should be investigated further. In this way, our visual approach can serve well as a wide-screen diagnostic tool for health monitoring in its current state.

For studies that do not require adverse event detection, the sensitivity of our method has added value. For instance, a drug or genetic manipulation that leads to slight (*∼*5%) change in mass can be detected accurately. Although a 5% change in mass may seem low, in human clinical trials for weight loss drugs, the number of participants who achieve at least a 5% decrease in mass is the primary result that is reported [29]. This level of sensitivity makes this a useful tool beyond health checks for preclinical animal studies.

While our models achieve good performance, they are not without error. These errors can be broken down into different types based on their source of introduction: equipment spatial resolution, limitations of our selected measurement, and observation interval. Since we image at a spatial resolution of 480x480 where the mouse occupies a small percentage (<1% of area) of the visible area, increasing the resolution of the image will reduce error. Our measurement is that of the silhouette of the mouse from a top-down viewing angle. This alone doesn’t capture the complex posture of the animal. Using calibrated depth cameras to provide a better understanding of the animal’s volume would reduce prediction error. Finally, observation interval limits how much information can be used to make a prediction. Ideally, predictions could be done with a couple frames. However, we observed that there is high variation in area across the course of an open field assay, as the mouse performs different behaviors. This variation necessitates longer observational intervals to ensure accurate body mass predictions.

We currently only test our system with single mice. This is a starting point for a convenient method of mass determination in animal production setting or certain studies. We envision such a system can be used to access mass in a changing table that is routinely used in mouse facilities. Future versions of this method can incorporate multianimal approaches where individual animals are segmented to determine its mass. For homecage settings, the camera may be much closer to the animal and will provide a much higher image resolutions. These presumably will create an easier task [30–32]. However, these conditions can also introduce perspective and lens distortion, which must be accounted for. In addition, bedding and occlusions may present challenges under these conditions. Future developments can start to handle these challenges. In addition, these methods can be extended to other organisms such as rats and even to non-human primates, where manual mass assessment is much harder.

## 5 Methods

### 5.1 Mice

All mice were obtained from The Jackson Laboratory production colonies. The mouse videos were obtained from a previously conducted strain survey of over 2400 mice from 62 strains, including at least 4 male and 4 for almost every strain [12, 33](See Table S1). We selected 2028 individual mice out of that survey, leaving out those with missing features in the data like age or arena recorded. Our mice vary in age from 7-26 weeks old, but most are 8-12 weeks old. Each mouse was weighed with a analytical laboratory scale (Ohaus) immediately prior to recording. The sampled body weights range from 9.1-54g with mean 24.7g.

### 5.2 Arena Recording Setup

All behavior methods and datasets have been previously described [12, 33, 34]. We have published detailed descriptions of our data acquisition methods, hardware, and software [15]. Briefly, all videos were recorded at 30 fps, have 8-bit monochrome depth, run for 55 minutes, and are saved as 480 *×* 480 px files. Each camera was mounted approximately 100 cm above each arena, with zoom settings tuned to 8px/cm. Variations in this zoom were normalized by the corner detection approach described in results [19]. We recorded individual mice in six near-identical open field arenas. The arenas are 52 *×* 52 *×* 23 cm, and built with white PVC plastic floors and gray PVC plastic walls. A white 2.54 cm chamfer was added to all inner edges for easier cleaning. Each arena was illuminated by an LED light producing 200-600 lux of light.

### 5.3 Statistical Analysis

To predict the weight of the mouse, we fit two simple linear regression models (M1, M2) and four multiple linear regression models (M3, M4, M5, M6) using video and animal-specific covariates. We describe our model as follows

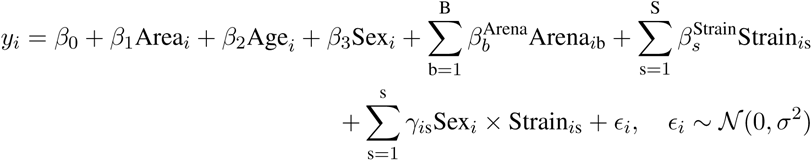

where y*_i_* denotes the weight of the mouse i, β*_p_*, p = 1, 2, 3, β_b_^Arena^, and β_s_^Strain^ are the regression coefficients associated with Area*_i_*, Age*_i_*, Sex*_i_*, Arena*_i_*, Strain*_i_* covariates for mouse i and γ*_i_* represents the interaction effect between Strain and Sex, ɛ*_i_* denotes the error term. The regression coefficients β and γ are estimated from the data using the least squares algorithm from the data so as to use the model to predict the weight of a new mouse. Further,

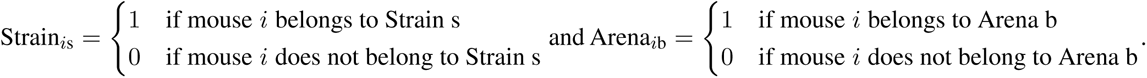

Each of our six models (M1-M6) can also be conveniently described as follows,

M1: Weight *∼* *A_px_*

M2: Weight *∼* *A_cm_*

M3: Weight *∼* *A_cm_* + Arena

M4: Weight *∼* *A_e_* + Arena

M5: Weight *∼* *A_e_* + Arena + Age + Sex

M6: Weight *∼* *A_e_* + Arena + Age + Sex + Strain + Sex * Strain

To assess the accuracy of our model predictions, we split the data randomly into two parts: train (70%) and test (30%). The test set served as an independent evaluation sample for the models’ predictive performance. We performed repeated cross-validation with 50 repeats to allow for a proper assessment of uncertainty in our test set results. The models were compared in terms of mean absolute error (MAE), mean absolute percent error (MAPE), root-mean-squared error (RMSE), and *R*^2^. These are defined as follows:

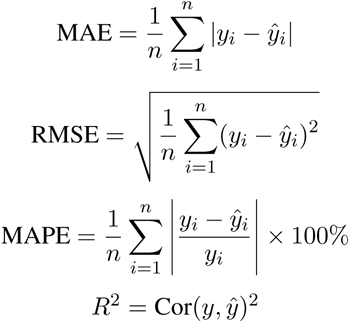

where *y*, *ŷ* are the true (observed) and predicted values from the model respectively.

We also performed Likelihood Ratio Tests (LRT) between each successive model that confirmed each next model provides a higher goodness of fit than the last, i.e. M2 provides significantly higher goodness of fit than M1 does, and so on. We fitted our models in R, performing statistical tests with the packages `lmtest` and `AICcmodavg`.

### 5.4 Data and Code availability

All raw video data will be shared on Harvard Data Verse upon acceptance of this manuscript. The segmentation neural network code and weights have been previously published and available [12]. All code is publicly available through the Kumar Lab GitHub (https://github.com/KumarLabJax/visual-mouse-weight/tree/v1.0.0).

## 6 Author contributions

M.G, B.G, and V.K. designed the experiments and analyzed the data. G.S. and M.G. carried out statistical modeling analysis. All authors wrote and edited the paper.

## 8 Acknowledgements

We thank Kumar Lab members for helpful advice and comments. We thank Kayla Dixon, a Colby-Lunder Fellow in the Kumar Lab, who initiated the project. This work was funded by The Jackson Laboratory Directors Innovation Fund, National Institute of Health DA051235, DA048634 (NIDA, V.K.), National Institute of Aging AG078530 (NIA, V.K.), National Institute for Neurological Disorders and Stroke NS078795 (NINDS, M.G.) and the C.C. Little Scholarship Fund (M.G.).

## 9 Supplementary Material

**Figure S1:**
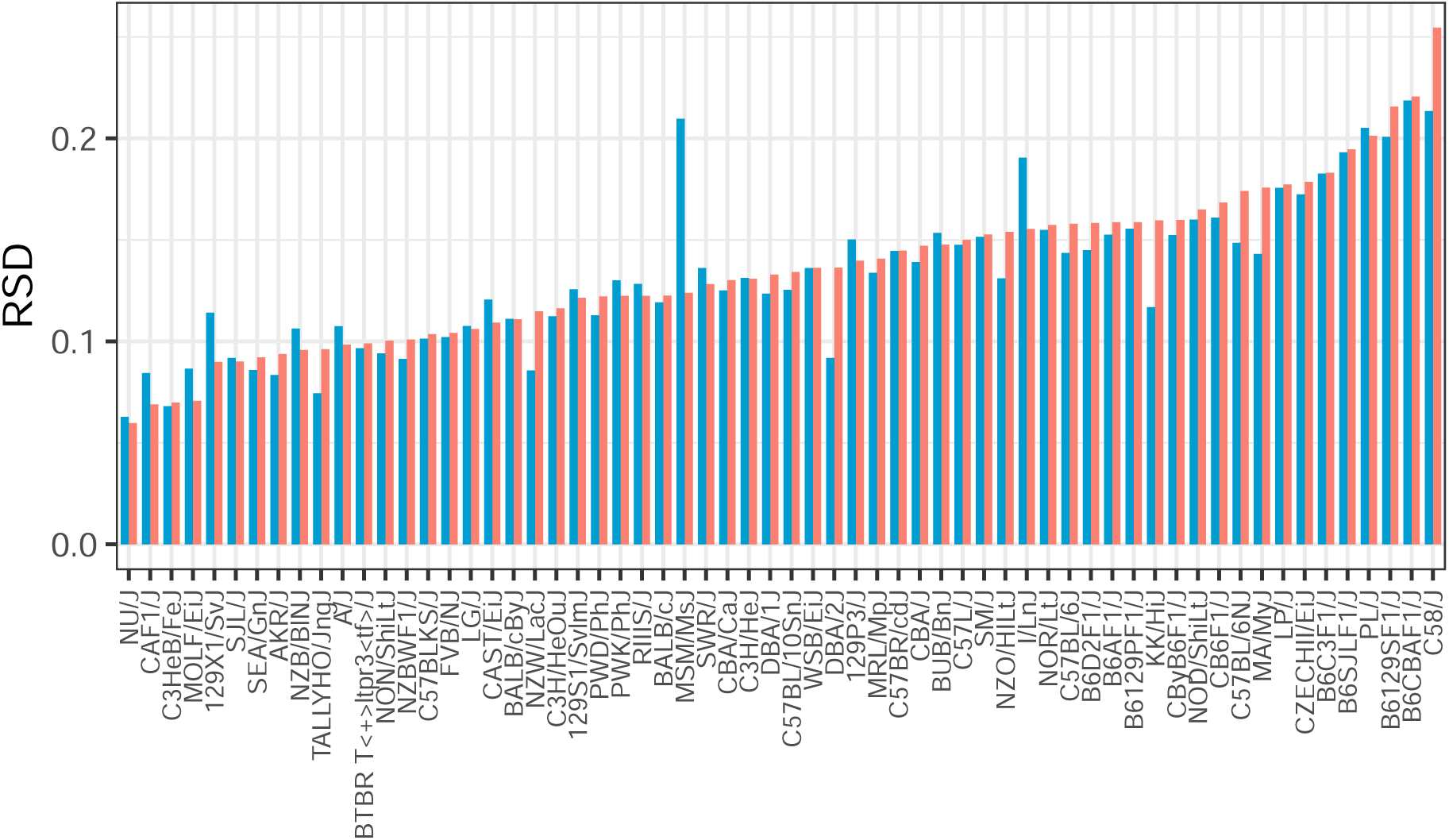
Relative Standard Deviations of Observed and Predicted Masses. Relative standard deviations (RSD) of observed mass (in orange) and the mass predicted by M6 (in blue, trained on the full dataset) for each of the 62 mouse strains considered in our dataset.

**Figure S2:**
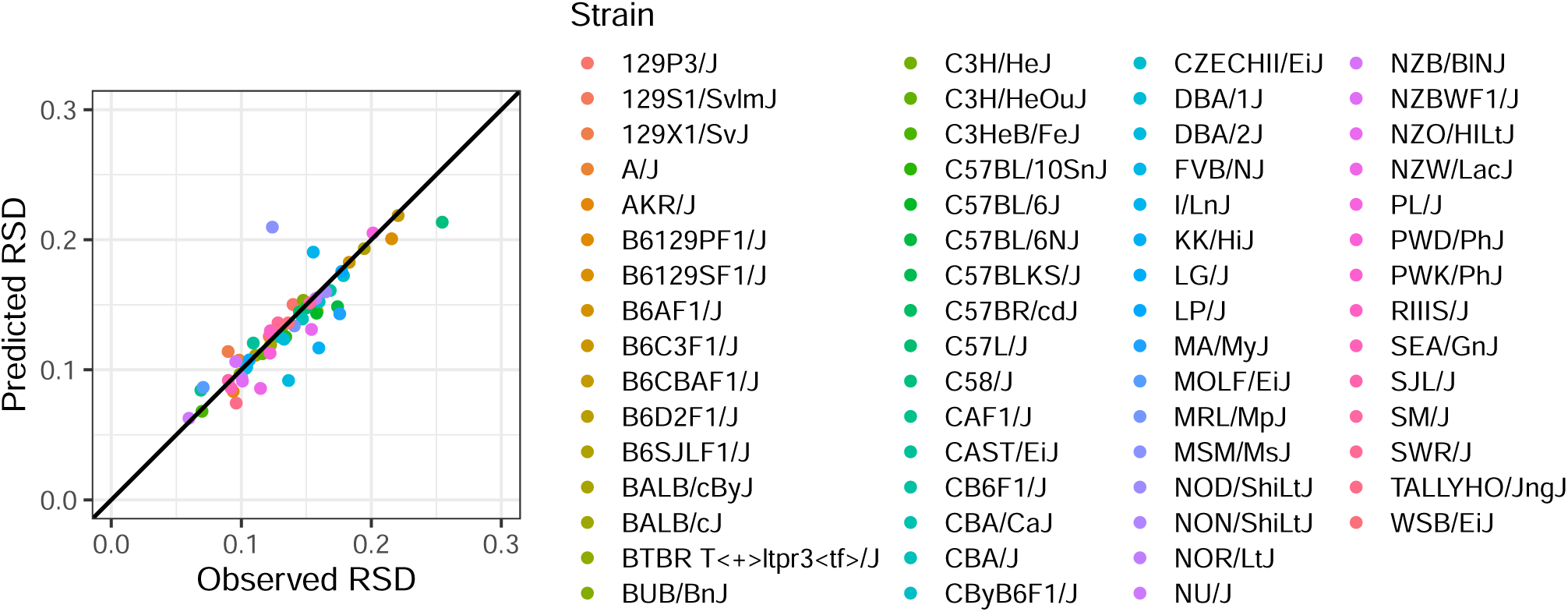
Observed and Predicted RSDs are Similar for M6 Across Strains. Relative standard deviations (RSD) of mean observed mass and mean predicted mass for each of the 62 mouse strains considered in our dataset. M6 is trained on the full dataset here. The black diagonal line is 45°, indicating exact correspondence between observed and predicted RSD.

**Table S1:**
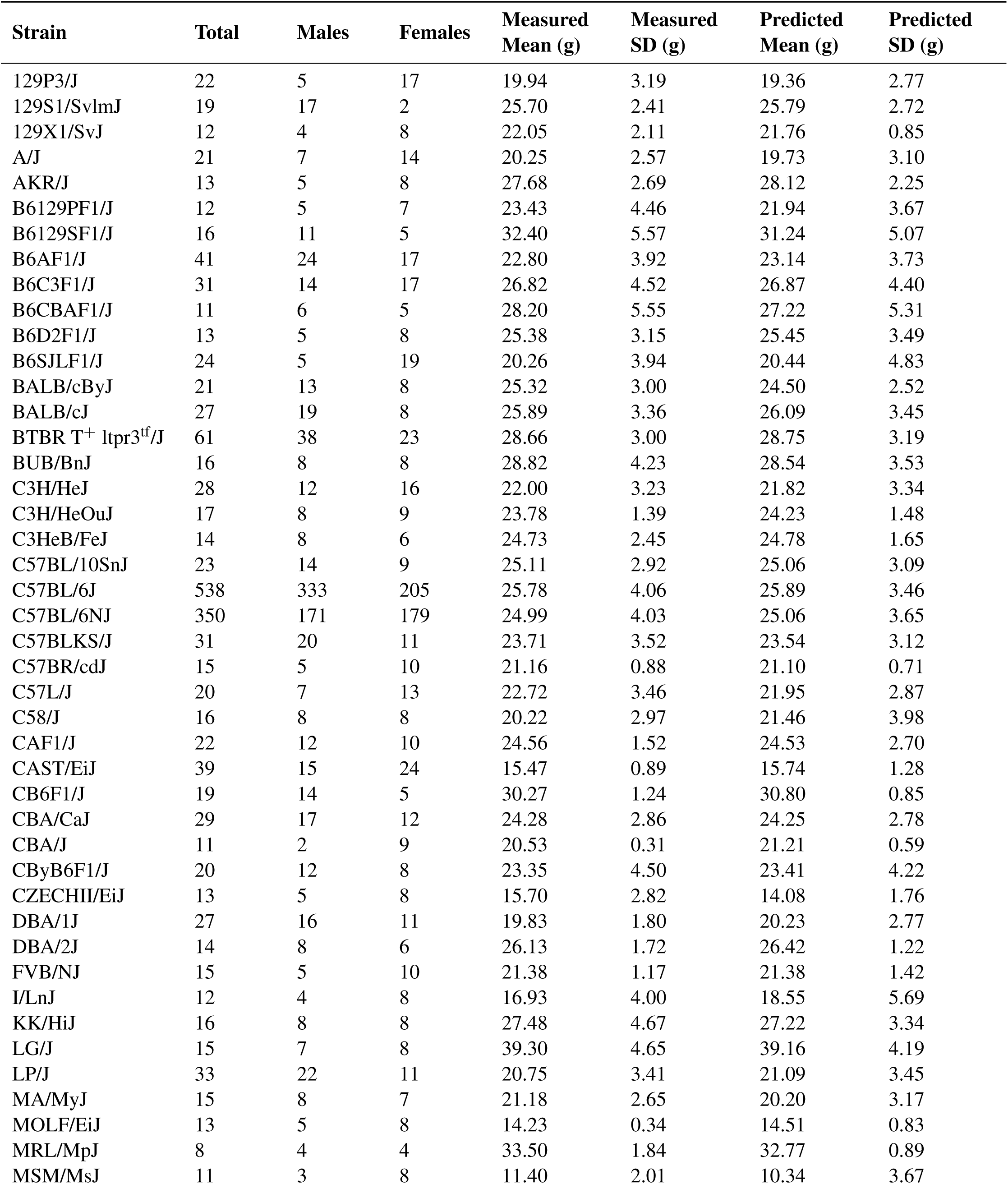
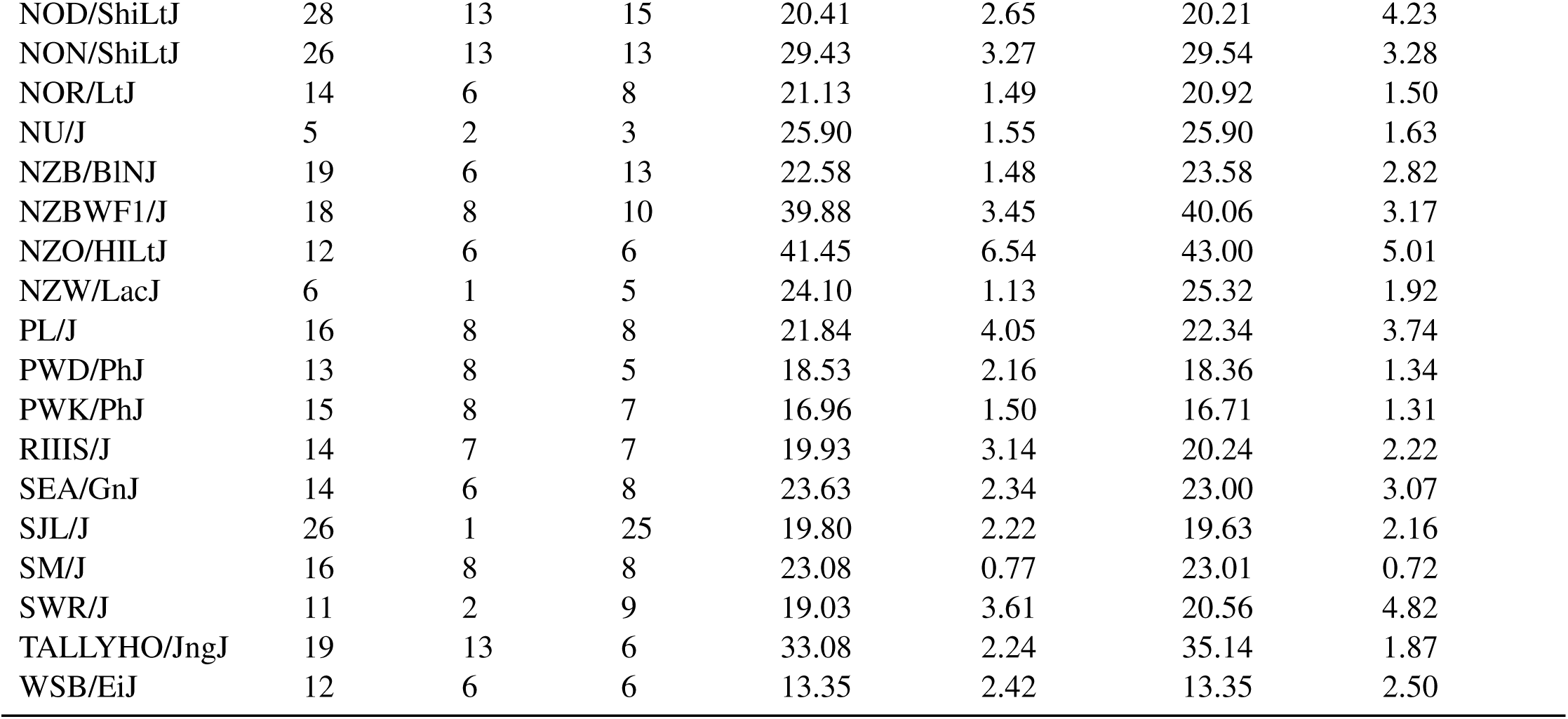
Full Model (M6) Performance by Strain. Mean measured and predicted masses reported for each strain, along with standard deviations. Number of male and female mice included in dataset from each strain reported.

